# The *Drosophila* TMEM184B ortholog Tmep ensures proper locomotion by restraining ectopic firing at the neuromuscular junction

**DOI:** 10.1101/2021.09.11.459917

**Authors:** Tiffany S. Cho, Eglė Beigaitė, Nathaniel E. Klein, Sean T. Sweeney, Martha R.C. Bhattacharya

## Abstract

TMEM184B is a putative seven-pass membrane protein that promotes axon degeneration after injury. *TMEM184B* mutation causes aberrant neuromuscular architecture and sensory and motor behavioral defects in mice. The mechanism through which TMEM184B causes neuromuscular defects is unknown. We employed *Drosophila melanogaster* to investigate the function of the *TMEM184B* ortholog, *Tmep* (CG12004) at the neuromuscular junction. We show that Tmep is required for full adult viability and efficient larval locomotion. *Tmep* mutant larvae have a reduced body contraction rate compared to controls, with stronger deficits in females. Surviving adult *Tmep* mutant females show “bang sensitivity,” a phenotype associated with epileptic seizures. In recordings from body wall muscles, *Tmep* mutants show substantial hyperexcitability, with many post-synaptic potentials fired in response to a single stimulation, consistent with a role for Tmep in restraining synaptic excitability. Neuromuscular junctions in *Tmep* mutants show modest structural defects and satellite boutons, which could also contribute to poor locomotor performance. Tmep is expressed in endosomes and synaptic vesicles within motor neurons, suggesting a possible role in synaptic membrane trafficking. Using RNAi knockdown, we show that Tmep is required in motor neurons for proper larval locomotion and excitability. Locomotor defects can be rescued by presynaptic knock-down of endoplasmic reticulum calcium channels or by reducing evoked release probability, suggesting that excess synaptic activity drives behavioral deficiencies. Our work establishes a critical function for the *TMEM184B* ortholog *Tmep* in the regulation of synaptic transmission and locomotor behavior.

## Introduction

Synaptic transmission at the neuromuscular junction (NMJ) requires precisely controlled anatomical and functional coordination between motor neuron terminals and their postsynaptic muscle partners. When action potentials arrive at synaptic terminals, calcium rise in the presynaptic terminal evokes the fusion of neurotransmitter-filled vesicles with the plasma membrane, triggering depolarization of muscle fibers. To fire a single action potential, the synaptic terminal must be able to restore ion levels to pre-stimulus levels quickly [1]. Mutations that fail in clearance of synaptic calcium [2], cause gain of function of calcium channels [3], or cause loss of the repolarizing function of potassium channels [4], show ectopic firing of synapses. This has deleterious effects on neural circuits and their resulting behavioral outputs.

The trafficking of presynaptic membrane proteins to their proper locations underlies the efficiency of synaptic transmission. Mutations that impair endocytic recycling of proteins leave presynaptic terminals ill prepared for repetitive firing events [5–9]. In addition, lipid dynamics at the synapse contribute to proper firing patterns [10–12]. Endosomes serve as sorting stations at the synapse and participate in the synaptic vesicle recycling process across species [5, 13–15]. The perturbation of vesicle recycling via the manipulation of membrane trafficking has been suggested as a therapeutic strategy for synaptic disorders [16].

The *TMEM184B* gene is a predicted 7-pass transmembrane protein that localizes to endosomal compartments, suggesting a possible role in intracellular membrane trafficking [17]. TMEM184b promotes efficient axon degeneration following nerve injury [17]. Paradoxically, *TMEM184B* mutations also cause substantially swollen nerve terminals, including at the NMJ. Mice deficient in TMEM184b show both sensory and coordination defects, indicating an important contribution to peripheral neuron function even in the absence of injury [17, 18]. Despite its clear contribution to synaptic architecture and behavior, its function at the synapse is not understood.

We sought to learn how TMEM184B influences synaptic structure and function using the *Drosophila* neuromuscular junction, a model glutamatergic synapse in an easily accessible system for structural, functional, and behavioral analysis. We have identified the *Drosophila* ortholog of *TMEM184B*, *CG12004*, which we named *Tmep* (transmembrane endosomal protein). Here we show that loss of Tmep, like TMEM184B, causes structural defects at the fly NMJ and impairs larval crawling ability. Flies deficient in Tmep show striking hyperexcitability at the NMJ, with many ectopic responses to a single stimulus. Despite broad expression, Tmep seems to control behavioral output via its motor neuron activity. Behavioral deficits caused by Tmep reduction can be rescued by depression of synaptic excitability through multiple mechanisms. Our data supports a presynaptic contribution of the TMEM184B ortholog Tmep in regulation of synaptic function and motor behavior.

## Results

### CG12004 (Tmep), the *Drosophila* ortholog of TMEM184b, is needed for full viability and locomotor function

To identify the closest ortholog to mouse and human TMEM184B, we used NCBI BLAST to select the most similar proteins in *Drosophila*. The CG12004 gene is 59% identical at the protein level to human TMEM184B, and reverse blast of CG12004 identifies TMEM184B as its closest relative. To understand which domains are most conserved, we mapped the aligned sequences onto the predicted transmembrane structure of CG12004. Two isoforms of CG12004 are predicted which differ only at their C-terminal (Fig 1a-b). The 7-pass transmembrane region, which has distant similarity to bile acid transporters, shows over 75% amino acid identity across fly, mouse, and human TMEM184B, indicating strong evolutionary conservation (Fig 1a). The third transmembrane domain is less strongly predicted than others but appears as a hydrophobic peak in all orthologs. Domains outside of the membrane are less well conserved, but a stretch of the membrane-proximal C-terminal shows high homology (67%, or 36 of 54 amino acids identical across all species), while the N-terminal is only 26% identical. Based on this analysis, CG12004 is the clear ortholog of mouse and human TMEM184B. Because TMEM184B localizes to endosomal compartments in cultured mammalian neurons [17], we have named the *Drosophila* isoform Tmep (<u>T</u>rans<u>m</u>embrane <u>e</u>ndosomal <u>p</u>rotein).

**Figure 1.**
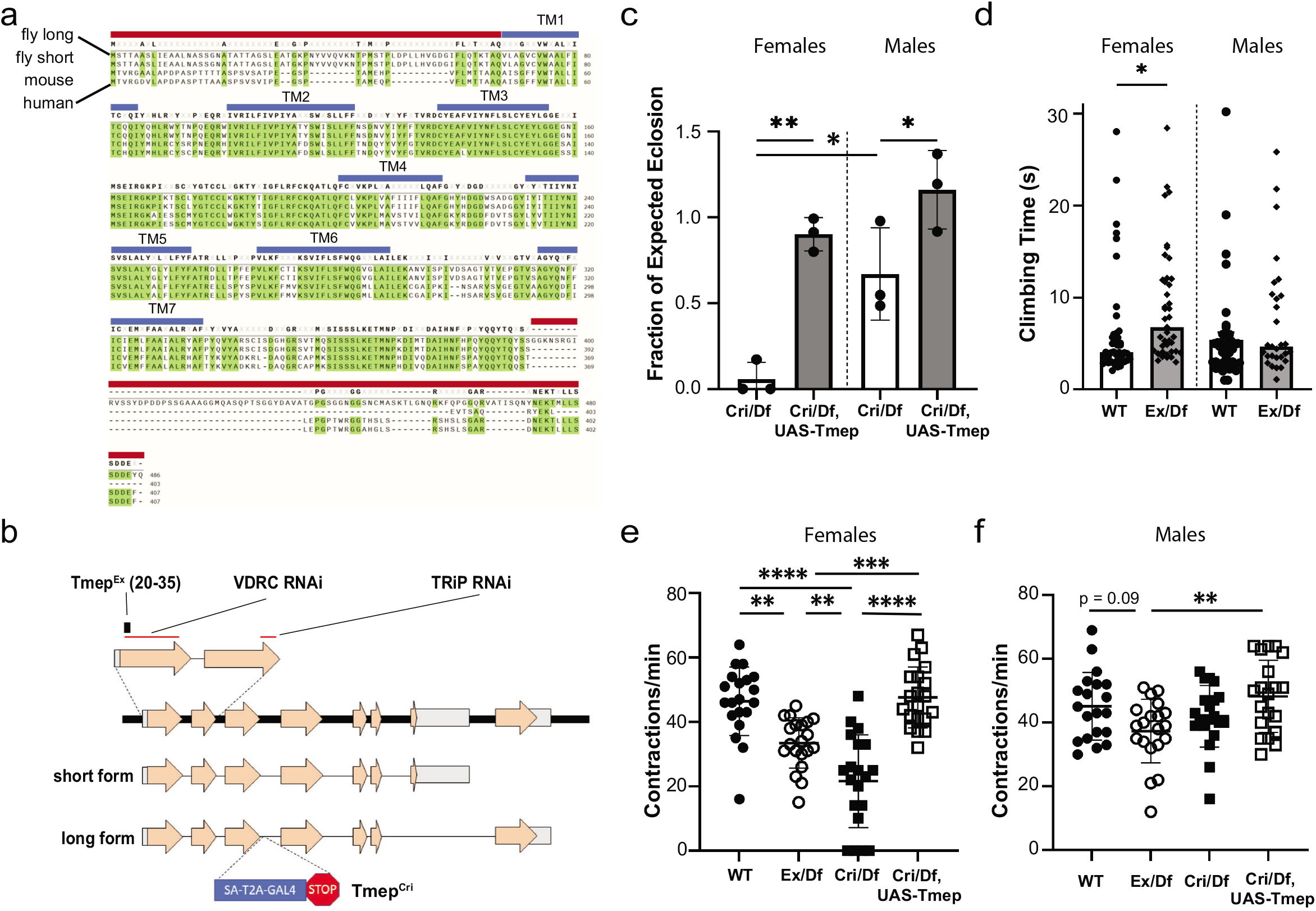
CG12004 (Tmep), the *Drosophila* ortholog of TMEM184b, is needed for full viability and locomotor function. **a**, Alignment of *Drosophila*, mouse, and human TMEM184B protein isoforms. Top to bottom: CG12004 long isoform, CG12004 short isoform, mouse TMEM184b isoform, human TMEM184b isoform. Transmembrane segments (blue lines) and sequences used for antibody generation (red lines) are shown. **b**, Schematic drawing of the CG12004 genomic region on Chromosome 3. Mutations and regions targeted by knockdown constructs used in this study are indicated. **c**, Rates of eclosion of *Tmep^Cri^*/Df (*Cri/Df*) in both sexes and rescue by UAS-Tmep (N-terminal mKate-tagged). P values: Male vs Female *Tmep^Cri^/Df*, p = 0.013; Male *Tmep^Cri^/Df* vs Rescue, p = 0.020; Female *Tmep^Cri^*/Df vs Rescue, p = 0.0006. N = 3 independent crosses with 5 male and 5 female parents in each cross. Statistical evaluation was done using two-tailed t-test within or across sex. **d**, bang sensitivity analysis of wild type and Tmep^*Ex/*^*Df* (*Ex/Df*) females and males. N = 43, 41, 42, 29 flies (left to right). For females, p = 0.045, unpaired t-test. **e-f**, larval contraction rate by genotype and sex (e shows females, f shows males). P values for females: wild type vs *Tmep^Ex^/Df*, p = 0.0017; wild type vs *Tmep^Cri^*/Df, p < 0.0001; *Tmep^Ex^/Df* vs *Tmep^Cri^/Df*, p = 0.005; *Tmep^Ex^/Df* vs rescue, p = 0.0005; *Tmep^Cri^/Df* vs Rescue, p < 0.0001; wild type vs rescue, p = 0.98. For males, all comparisons were p > 0.05 except *Tmep^Ex^/Df* vs rescue, where p = 0.008. N = 20 for all genotypes. Statistical significance was calculated by one-way ANOVA with Tukey’s multiple comparison correction.

We used homologous recombination to create a Tmep mutant allele, *Tmep^Ex^*, with a 16-base excision (bases 20-35 in the open reading frame) that is predicted to create a nonsense mutation 7 amino acids from the N-terminal, affecting both the short and long Tmep isoforms (Fig 1b). We also obtained a second allele (*Tmep^Cri^*) in which Tmep was disrupted by a CRISPR-Mediated Integration Cassette (CRIMIC) cassette containing a splice acceptor, *GAL4* sequence, and a “stop” signal [19]. Both *Tmep^Ex^* and *Tmep^Cri^* disrupt all four predicted *Tmep* transcripts. We used these two alleles in transheterozygous combination with a deficiency, *Df(3L)Exel6087*, that uncovered the Tmep locus, as well as two RNAi constructs targeting different conserved sequences (Fig 1b), in our subsequent studies.

When setting up transheterozygous crosses to study *Tmep* mutant flies, we observed that *Tmep^Cri^*/Df flies had severely impaired rates of eclosion compared to non-mutants in the same vial (Fig. 1c). *Tmep*-associated lethality was greater in females than in males. This lethality is often observed prior to wandering third-instar stage, as relatively few mutant larvae with this allele combination can be found (not shown). In both sexes, Tmep re-expression driven by the *CRIMIC-GAL4* (here used as both a mutant allele and a rescue driver in the Tmep expression pattern) was sufficient to rescue eclosion rates to wild type levels (Fig. 1c). Due to the severe lethality of *Tmep^Cri^*/Df, we analyzed *Tmep^Ex^/Df* in subsequent experiments.

Because *Tmem184b* mutant mice show sensory and motor defects [17, 18], we wanted to know if *Tmep* mutant flies would show similar phenotypes. First, we evaluated if flies showed “bang sensitivity,” an assay used to evaluate seizure and paralytic behavior [20]. *Tmep* mutant females showed slower climbing time after banging, while males were not affected (Fig 1d). In larvae, we assessed how *Tmep* reduction impacted locomotor behavior using a simple crawling assay. In females, both *Tmep^Ex^* and *Tmep^Cri^*, in trans to a deficiency, showed reduced body contraction (crawling) rates compared to controls (Fig. 1e), with *Tmep^Cri^* having stronger effects. Re-expression of *Tmep* driven by *CRIMIC-GAL4* was sufficient to rescue crawling behavior in females (Fig. 1e). In males, while behaviors trended similarly to females, none of the *Tmep* mutant genotypes were significantly different from wild type (Fig 1f). Taken together, our data show that Tmep expression is critical for viability and behavior, with effects of Tmep reduction more deleterious in females. We do not yet know the basis for the sexual dimorphism in *Tmep* mutants. In subsequent experiments, we separated animals by sex to fully capture the phenotypes caused by Tmep reduction.

### Tmep restrains ectopic firing at neuromuscular junctions

Bang sensitivity can be caused by mutations in ion channels and other pathways that affect synaptic transmission and nerve excitability [20]. In mice, *Tmem184b* mutation affects performance on broad sensorimotor tests, but does not alter conduction velocity or action potential amplitude in sensory or motor nerves [17]. Therefore, we hypothesized that Tmep loss may impact synaptic transmission rather than action potential propagation, leading to behavioral defects. To investigate whether Tmep is required for proper synaptic transmission, we recorded both spontaneous (mini excitatory junctional potential, mEJP) and evoked (EJP) activity in 3^rd^ instar larvae. In wild type larvae, a single evoked action potential triggers a single EJP spike. However, when action potentials were triggered in *Tmep^Ex^* /Df larvae, we saw a striking hyperexcitability phenotype (Fig 2a). Both males and females commonly responded more than once to a stimulus, and occasionally a single stimulus induced over 20 EJPs over the 5 second post-stimulus period (Fig 2b). While the first EJP did not vary in amplitude between wild type and mutants (Fig 2c), all second (ectopic) EJPs had reduced amplitudes in comparison to the first response (Fig 2d-e), though this difference was not statistically significant. Ectopic EJPs occurred relatively rapidly after the stimulus, with spacing between ectopic responses lengthening slightly by the 5^th^ response (Fig 2f).

**Figure 2.**
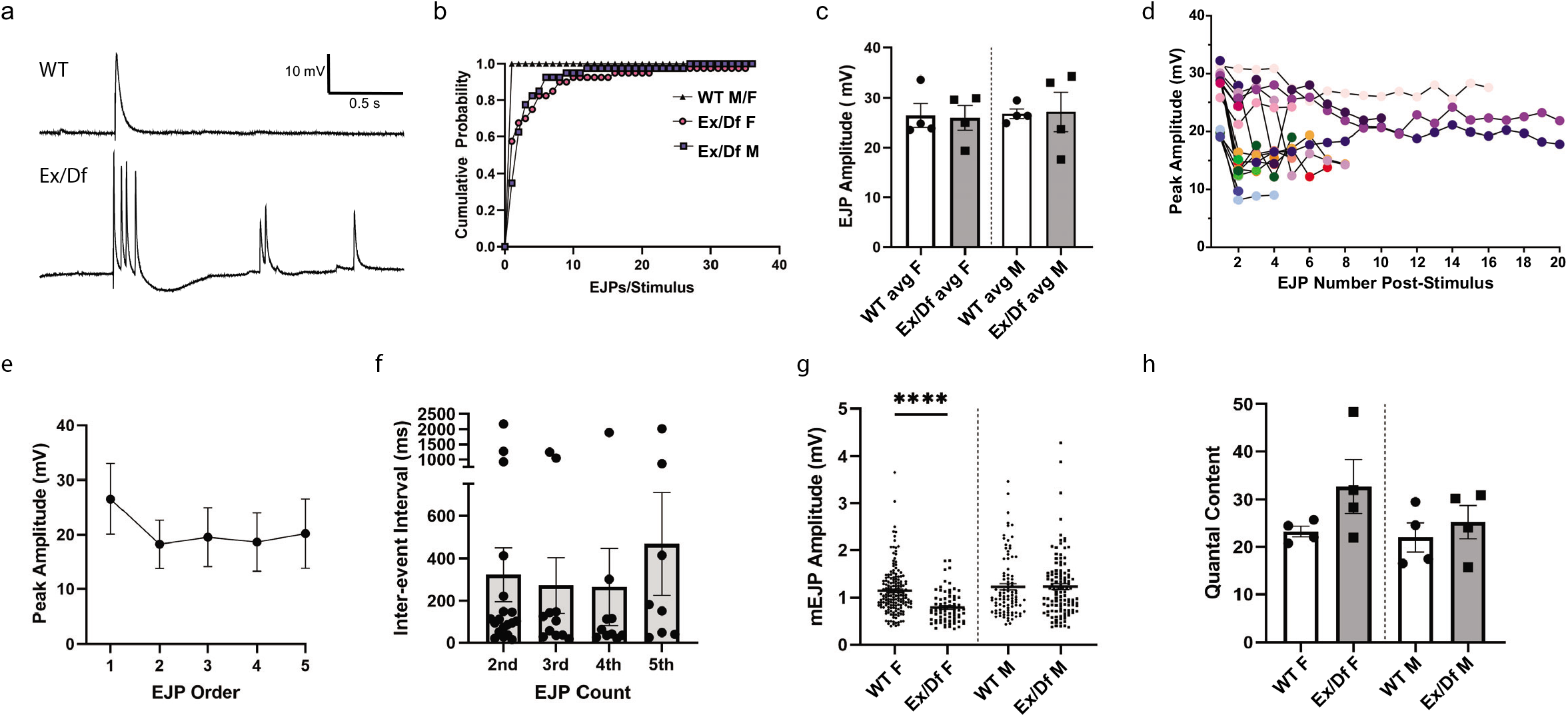
Tmep is required for proper synaptic transmission. **a**, Representative examples of evoked junctional potentials (EJPs) evoked by stimulation of wild type (WT) or *Tmep^Ex^/Df* (*Ex/Df*) female neuromuscular junctions (1 ms, 3.3 mV). **b-e**, Analysis of EJPs from N=4 larvae/genotype and n=10 stimuli/larvae. All EJP analysis is from the same animals. **b**, Cumulative probability of single and multiple EJPs following stimulation in wild type and *Ex/Df* larvae (over the 5 sec post-stimulation), separated by sex. All recordings were done in 0.4 mM Ca^2+^ HL3.1. **c**, EJP amplitudes of the first response recorded following each stimulus. **d**, Peak amplitudes of EJPs per stimulus. 17 of 40 EJPs measured in 4 *Ex/Df* females showed multiple peaks and are displayed on the graph. **e**, Relative sizes of multi-EJP responses to a single stimulus from female *Ex/Df* larvae. N=17, 17, 13, 12, and 10 larvae showed at least 5 responses and were included in analysis. No significant differences were measured (p > 0.05 for all comparisons). **f**, Inter-event interval from females shown in e. Only the first five inter-event intervals were analyzed, and each point represents the interval from the previous spike (i.e. 2^nd^ is from spike 1 to spike 2). No significant differences were observed (p > 0.05, one-way ANOVA). **g**, mEJP amplitudes from wild type and *Ex/Df* larvae of each sex. N=4 larvae per group. Fifteen seconds of baseline recording were analyzed per animal. T-tests within each sex were used to evaluate statistical significance (females, p <0.0001; males, p = 0.98). **h**, Quantal content calculation from wild type and *Ex/Df* larvae of each sex. Calculation was done per animal by dividing the first EJP (in **c**) by the average mEJP of that larva (in **g**). No significant differences were measured (p > 0.05 for all comparisons).

When examining spontaneous events (mEJPs), we found a reduction in the amplitudes of mEJPs in female, but not male, *Tmep* mutants (Fig 2g). However, this change did not affect quantal content, suggesting similar numbers of presynaptic vesicles available for release (Fig 2h). These data show that Tmep normally restrains evoked synaptic responses. Taken together with the behavioral data, our recordings support a model where, in the absence of Tmep, excessive firing at NMJs impairs locomotor ability.

### Tmep localizes to endosomes and synaptic vesicles in motor neurons

TMEM184B is expressed in peripheral neurons (motor and sensory) as well as in the central nervous system in the mouse, starting from early development [18]. To understand why *Tmep* mutant larvae show hyperexcitability, we first wanted to establish its overall expression pattern at the larval stage. We developed a polyclonal antibody to *Drosophila* Tmep (regions used as antigens shown by red lines in Fig 1a). Tmep is expressed in a punctate pattern in the neurons of the ventral nerve cord (Fig. 3a). Importantly, we see very low signal in *Tmep^Ex^/Df* larvae, indicating that our antibody is specific. Tmep is also found at low levels at the NMJ (Fig. 3b).

**Figure 3.**
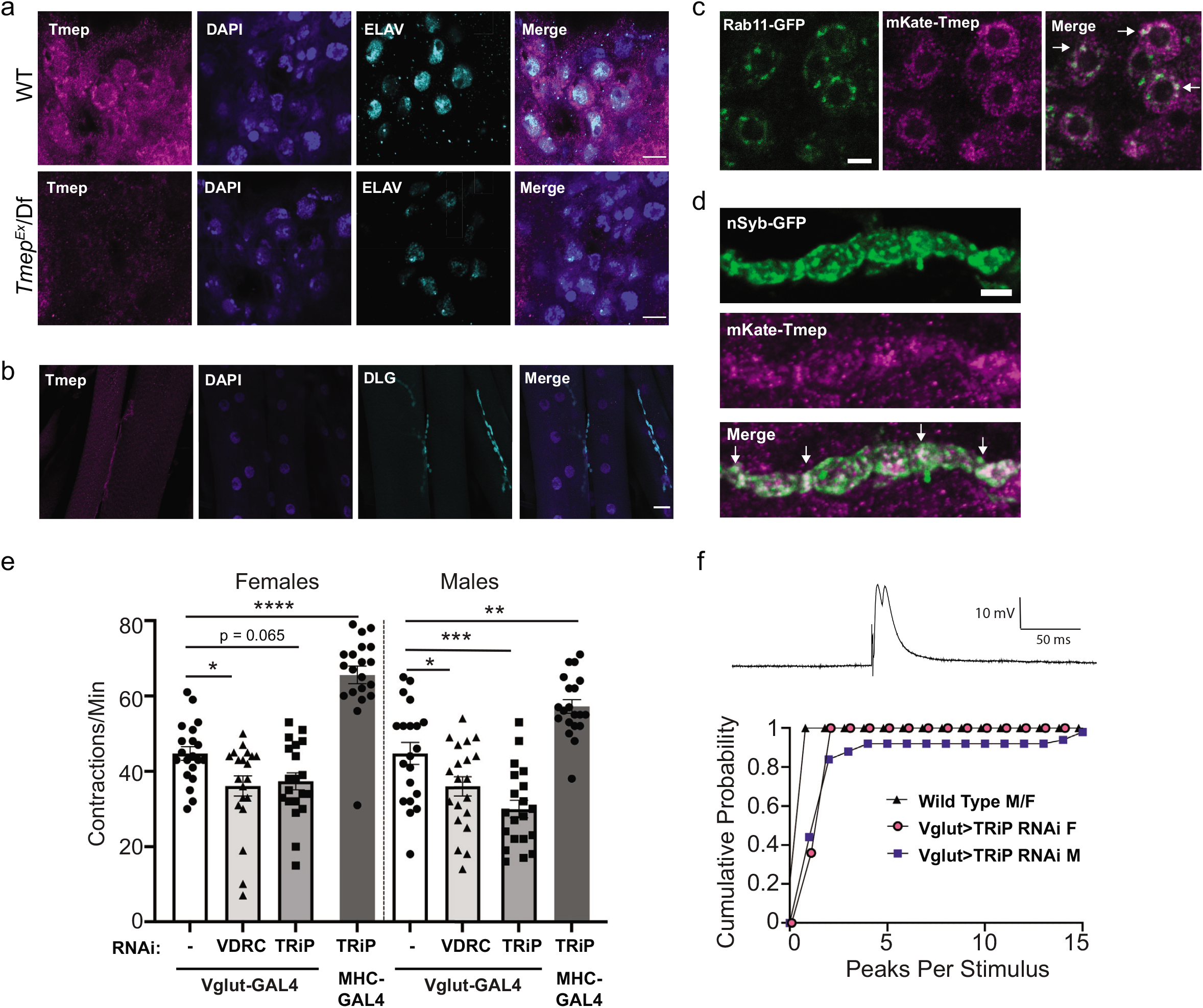
Tmep localizes to neuronal endosomes and is required presynaptically for evoked responses and locomotion. **a**, Antibody to *Drosophila* Tmep shows neuronal staining in the larval ventral nerve cord. Magenta, Tmep; Blue, DAPI; Teal, Elav (neuronal marker). Limited staining is observed in Tmep mutants (*Tmep^Ex^/Df*). Scale bar = 5 μm. **b**, Tmep protein levels at the neuromuscular junction. Magenta, Tmep; Blue, DAPI; Teal, *discs large* (DLG) showing the postsynaptic area. Scale bar = 30 μm. **c,** Tmep shows partial colocalization with Rab11-positive endosomes. Motor neuron expression of mKate-Tmep (magenta) and Rab11-GFP (green) is driven by *BG380-GAL4*. Image is a single confocal slice of cell bodies in the ventral neve cord; arrows in the merged image show areas of partial colocalization. Scale bar = 5 μm. **d**, Tmep shows partial colocalization with synaptic vesicles at the larval NMJ. Image is a single confocal slice. Nsyb-GFP and mKate were expressed as in c. Scale bar = 5 μm. **e,** Crawling analysis of larvae in which Tmep is knocked down in motor neurons (*Vglut-GAL4*) or muscle (*MHC-GAL4*) with one of two different RNAi constructs (sequence targets are shown in Figure 1). N=20 animals per genotype and sex. One-way ANOVA with Dunnett’s post-hoc test within sex; p = 0.025 (Vglut F alone vs VDRC F), p = 0.065 (Vglut F alone vs TRiP F), p = <0.0001 (Vglut F alone vs MHC> TRiP F), p = 0.035 (Vglut M alone vs TRiP M), p = 0.0002 (Vglut M alone vs TRiP M), p = 0.0016 (Vglut M alone vs MHC> TRiP). **f**, representative trace (above) and cumulative probability (below) of multi-peak events in Tmep knockdown (TRiP) larvae. Note that controls (black triangles) do not show any multi-peak responses. N=5 larvae and 10 stimuli/larvae per genotype.

We wanted to evaluate if, as in mice, Tmep is found on Rab11-positive recycling endosomes. Using an mKate-tagged Tmep, we found a similar punctate, vesicular pattern as seen with the Tmep antibody. These small Tmep-positive puncta showed partial overlap with larger, Rab11-positive areas in motor neuron cell bodies (Fig 3c). In addition to direct overlap, Tmep puncta were often adjacent to Rab11-positive puncta, suggesting that Tmep-positive compartments may at least transiently intersect with Rab11-positive endosomes. We also observed substantial overlap between mKate-Tmep and GFP-tagged neuronal Synaptobrevin (n-Syb) (Fig 3d), which is localized to synaptic vesicles as well as synaptic endosomal compartments [21]. Taken together, our data indicate that Tmep is primarily expressed on intracellular membranes, including endosomes and synaptic vesicles. This localization is consistent with a role in synaptic transmission.

We also took advantage of the *CRIMIC-GAL4* insertion to express nuclear-localized RFP in cells expressing endogenous Tmep. Tmep is expressed in muscle and motor neurons, but not in glia in the larval brain (Online Resource 1a-b). We also observe Tmep expression in the gut and fat bodies (Online Resource 1c-d). These data indicate that Tmep is broadly expressed in *Drosophila* larvae.

**Online Resource 1.**
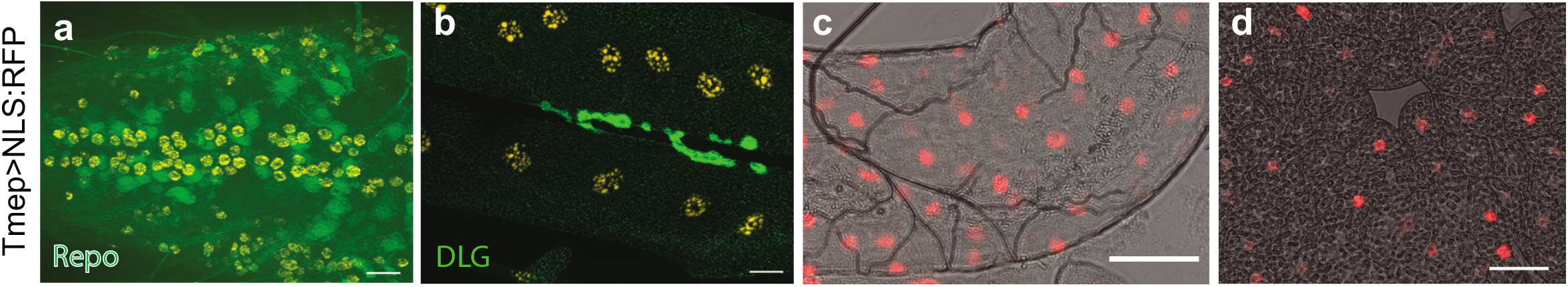
Tmep is expressed broadly throughout the larvae. RedStinger:NLS (red and orange) shows nuclei of cells in which endogenous Tmep is expressed. **a**, Ventral nerve cord. Green is Repo (glial nuclei). **b**, Neuromuscular junction. Green is *discs large* (DLG). **c**, Gut expression. **d**, Fat body expression. For all figures, scale bars = 20 μm.

### Tmep is required presynaptically for neuromuscular phenotypes

Tmep is expressed both by motor neurons and muscle. To establish where it contributes to locomotor function and physiology, we used tissue-specific RNA interference. When we knock down Tmep with two separate RNAi’s in motor neurons (*Vglut-GAL4*) we observe a reduction in larval crawling rate (Fig 3e) similar to that seen in genetic mutants (Fig 1e). The effect of knockdown is observed in both male and female larvae. Knockdown of Tmep in muscle (using *MHC-GAL4*) does not impair crawling; surprisingly, it increases contraction rate, indicating a possible feedback mechanism at play (Fig 3e). When we recorded EJPs from larvae with presynaptic Tmep knockdown, we observed a subtle but persistent multi-peak EJP in response to a large fraction of stimuli (Fig 3f). Fully independent ectopic EJPs were rare but did occur in some larvae. These data support a primarily presynaptic function for Tmep in controlling synaptic transmission and motor function.

### Tmep mutant larvae show morphological defects at the neuromuscular junction

*Tmep* locomotor and electrophysiological phenotypes in *Drosophila* larvae could be caused by morphological changes at the synapse, alterations in synaptic signaling pathways, or a combination of both. To investigate if changes in NMJ morphology could contribute to phenotypes, we examined terminal morphology with immunohistochemistry. In mice, disruption of TMEM184B expression causes swellings in the NMJ [17]. We observed statistically significant increases in nerve terminal branching in *Tmep* mutant female larvae (Fig 4a-b). In *Tmep* mutant presynaptic terminals, we observed numerous satellite boutons (Fig 4c-e). Satellite or fragmented bouton architecture is also observed at the ultrastructural level (Fig 4d). Our results are consistent with those made at the mouse NMJ and suggest that morphological changes could contribute to *Tmep* phenotypes [17].

**Figure 4.**
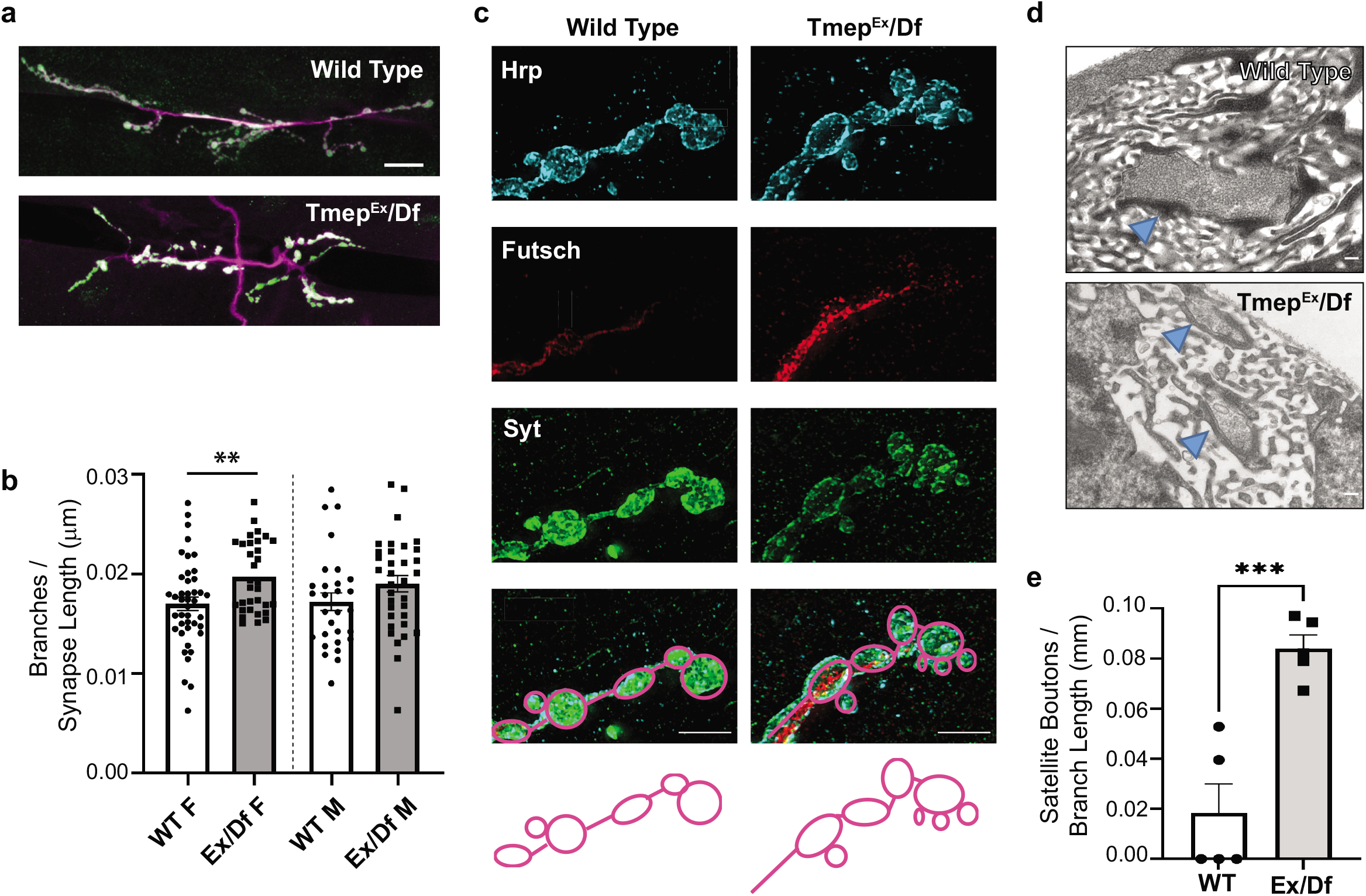
Tmep mutant larvae show extra branches and boutons at the neuromuscular junction. All images show wandering female third instar larvae. **a**, Representative images of muscle 6/7 NMJs stained with antibodies to Synaptotagmin (Syt, green) and HRP (magenta) illustrating branching and bouton phenotypes. Scale bar = 20 μm. **b**, Quantification of branches in wild type or *Ex/Df* larvae. Branch count is normalized to the total synapse length (sum of all branches). **c**, Synaptic terminal boutons in wild type and *Ex/Df* showing an example of satellite boutons at the mutant NMJ. HRP (blue), Futsch (red), and Synaptotagmin (Syt, green) are shown. Scale bar = 5 μm. Drawings illustrate the bouton structure in the above images. **d**, Transmission electron micrographs showing examples of synapses in wild type and *Ex/Df* larvae. Scale bars = 1 nm. Arrows indicate synaptic sites and show multiple smaller terminals in *Ex/Df* larvae, which could be satellite boutons or fragmented synaptic endings. **e**, Satellite bouton quantification normalized to the length of the branch upon which the boutons are located. N = 5 female larvae per genotype. p = 0.0009 using unpaired t-test.

### Larval crawling deficiencies are restored by reduction of presynaptic excitability

Ectopic EJPs are seen in *Drosophila* mutants that impair the clearance of calcium from presynaptic terminals [2, 3]. The endoplasmic reticulum and its stored calcium positively influence synaptic transmission [22, 23]. Therefore, we asked if reducing cytoplasmic calcium levels in motor neurons could improve Tmep-knockdown locomotor behavior. We first blocked store-operated calcium release in two ways, using RNAi to either the inositol triphosphate (IP_3_) receptor (ITPR) or the ryanodine receptor (RyR). While the individual RNAi’s to each receptor did not change baseline locomotion, both ITPR and RyR knockdown were able to restore normal locomotor activity in the background of Tmep RNAi (Fig 5a-b). This fits with the emerging roles for presynaptic ER signaling in the regulation of synaptic transmission [24–26].

**Figure 5.**
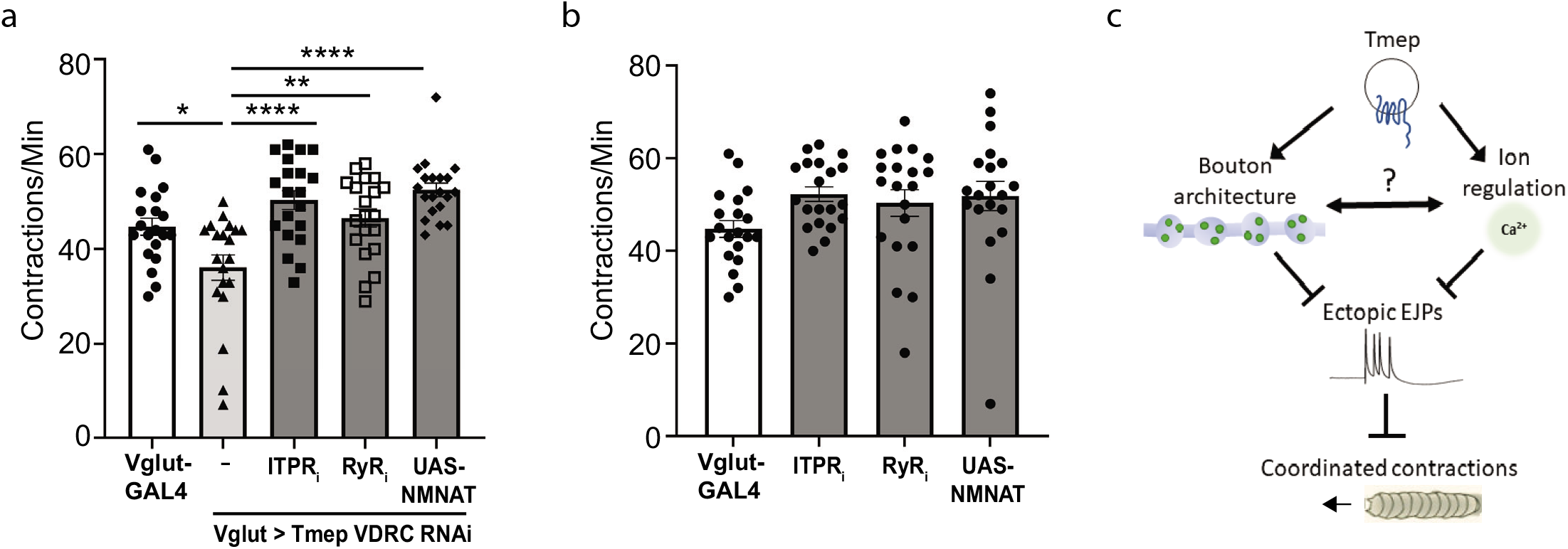
Larval crawling deficiencies are restored by reduction of presynaptic excitability. **a**, Crawling deficiency of Tmep knockdown larvae can be restored by knockdown of IPTR, RyR, or by over-expression of NMNAT. Female data is shown. One-way ANOVA with Sidak’s multiple comparison correction was used to calculate p values as follows: p = 0.014 (Vglut vs Tmep RNAi), p = 0.0014 (Tmep RNAi vs Tmep/RyR), p < 0.0001 (Tmep RNAi vs Tmepi/ITPRi or NMNAT). **b**, Individual RNAi expression of IPTR, RyR, or UAS-NMNAT do not change baseline contraction rates (for all, p>0.05). Female data is shown. **c**, Model for Tmep influence on synaptic activity and locomotion.

We also dampened presynaptic activity using an orthogonal strategy. The NAD+ synthetic enzyme NMNAT restrains evoked synaptic transmission in mutants where synaptic activity is enhanced, such as *highwire* (*hiw*) [27]. In addition, over-expression of NMNAT in a wild type background dampens evoked release from motor neurons [27]. Indeed, we found that by driving additional NMNAT expression in motor neurons in a Tmep RNAi background, crawling behavior was restored to control levels (Fig 5a-b). While our data does not definitively prove that NMNAT is directly regulated by Tmep (and indeed, NMNAT has many functions including maintenance of axon viability [28–30]), it supports the hypothesis that presynaptic hyperexcitability is the cause of crawling deficiencies in Tmep mutant and knockdown larvae.

Our data support a model in which Tmep, working in an endosomal or vesicular compartment, influences both bouton architecture and ion regulation at the synapse (Fig 5c). In *Tmep* mutants, dysregulation of these processes leads to ectopic evoked potentials, which reduces the coordination of muscle contractions necessary for larval locomotion.

## Discussion

In this work we have identified a role for the TMEM184b ortholog Tmep in promoting proper neuromuscular junction architecture as well as restraining ectopic firing at the glutamatergic synapses of the *Drosophila* NMJ. Excessive synaptic activity can result in epileptic syndromes. In epilepsy, excitatory-inhibitory balance is often disrupted in favor of over-excitation [31, 32]. Our data indicates that Tmep works presynaptically to ensure both the architecture of the NMJ and the proper responses to nerve stimulation. Interestingly, we have recently identified human patient variants in *Tmem184b*, one of which causes myoclonic epilepsy as well as other neurological syndromes (K. Chapman and S. Berger, personal communication). We hope to delve further into the links between Tmem184b structure, function, and excitability in the future by analyzing the effect of known patient variants on the phenotypes we describe in this manuscript.

While we do not yet precisely know the nature of Tmep’s contribution to synaptic hyperexcitability, we speculate that it controls a membrane trafficking function at the presynaptic terminal. We found that Tmep, like mouse TMEM184B [17], is localized to endosomal compartments. Interestingly, we see significant co-localization of Tmep with n-Syb, a protein present on both endosomes and synaptic vesicles. Loss of n-Syb causes the accumulation of endosomes [21], a phenotype reminiscent of the endolysosomal accumulations seen in mouse *TMEM184B* mutants. Along with the role of the yeast ortholog Hfl1 in vacuolar degradation [33, 34], these data implicate Tmep in endolysosomal progression.

Tmep would thus join a group of endosomal and membrane trafficking effectors whose mutation impairs synaptic transmission. For example, Rab5 promotes efficient release of synaptic vesicles by ensuring proper membrane exchange between these vesicles and synaptically localized endosomes [13, 35]. In addition, mutations in the NPC1 gene that cause Niemann-Pick type C1, a fatal neurodegenerative disease caused by faulty lysosomal-ER cholesterol transfer, also show ectopic responses to presynaptic stimulation [12]. These studies underscore the need to regulate membrane trafficking to ensure proper responses to stimulation. A priority for future work should be to investigate how Tmep or its mammalian counterparts influence essential functions of endosomes, lysosomes, and synaptic vesicles at neuromuscular junctions.

We observed that knocking down store-operated calcium channels was able to rescue *Tmep* mutant crawling behavior, indicating that Tmep, or potentially an associated endosomal compartment, may have a role in calcium dynamics within the synapse. ITPR is expressed on the endoplasmic reticulum and plays specialized roles in loading calcium into compartments such as mitochondria [36] and lysosomes [37]. The accumulation of lysosomes in mouse TMEM184b mutants [17] is consistent with an alteration that disrupts their function, perhaps one that alters lysosomal calcium storage. Loss of function of Tmep could also perturb plasma membrane calcium channel distribution via an endosomal function, in a similar manner to the role of Rab11 in N-type calcium channel recycling [38]. Another possibility is that Tmep assists in establishing or maintaining membrane contact sites to ensure proper calcium handling in membrane-bound organelles [39, 40]. Any disruption of calcium levels at the synapse would be predicted to cause changes in excitability, leading to the phenotypes we observe in *Tmep* mutants.

In summary, we have established a *Drosophila* model of *Tmem184b* associated nervous system phenotypes and have discovered a role for the fly ortholog, Tmep, in synaptic excitability and behavior. The evolutionary conservation between fly *Tmep* and human *TMEM184B* is strong, indicating a critical function across eukaryotic organisms. Our work establishes a system in which to study how Tmep/TMEM184B influences synaptic transmission and may contribute to human nervous system disorders.

## Acknowledgements

The authors would like to thank all members of the Bhattacharya lab for thoughtful comments on the manuscript. We also thank Dr. Konrad Zinsmaier for sharing his electrophysiology equipment as well as his expertise for our work. We thank Dr. Kimberly Chapman and Dr. Seth Berger for communicating unpublished information. We thank the York Protein Production Facility, the York Imaging and Cytometry Facility, and the University of Arizona Marley Microscopy Core Facility for their assistance.

## Declarations

### Availability of data and material

Flies and plasmids created during this study will be available from the authors or deposited at the Bloomington Drosophila Stock Center (BDSC) or at Addgene.

### Code availability

Not applicable.

### Authors’ contributions

S.T.S. and M.R.C.B. proposed the research and assisted in experimental design. T.S.C., E.B., and N.E.K. designed and performed experiments. All three first authors (indicated with *) made significant discoveries leading to this manuscript, resulting in all three being listed as equal first-authors. T.S.C. and M.R.C.B. wrote the manuscript, with editing by E.B. and S.T.S.

### Ethics approval

N/A

### Consent for publication

All authors have had the opportunity to review this manuscript prior to submission, and all give their approval for publication of this work.

## Materials and Methods

### Fly stocks and husbandry

Flies were raised at 25°C, and all experiments were performed at 22°C. Fly stocks were from either the Bloomington Drosophila Stock Center (BDSC) or the Vienna Drosophila RNAi Consortium (VDRC) as follows: *Vglut*-GAL4 (BDSC # 24635), *Df(3L)Exel6087* (BDSC #7566), *Tmep^Cri^* (BDSC #91319), *MHC82*-GAL4 [41], UAS-CG12004 RNAi (TRP #57429), UAS-CG12004 RNAi (VDRC #101732), UAS-ITPR RNAi (BDSC #51795), UAS-RyR RNAi (BDSC #28919), UAS-NMNAT (BDSC # 39699), and wild type (Canton S or *w^1118^*). *Tmep^Ex^* was made by InDroso Functional Genomics (Rennes, France) by homologous recombination to create a 16-base deletion in the first coding exon of CG12004. UAS-mKate:CG12004 (long form) was created by cloning into the pBID vector, which was a gift from Brian McCabe (Addgene plasmid # 35190; http://n2t.net/addgene:35190; RRID:Addgene_35190). Injections were done at Bestgene (Chino Hills, CA).

### Genome Alignment

Sequences were downloaded from NCBI as follows: human TMEM184b (isoform a, NP_036396.2), mouse TMEM184b (isoform 1, NP_766196.1). Fly orthologs were derived from Flybase (CG12004-RD is the long form, CG12004-RA is the short form). Alignments were done in Snapgene using ClustalW. Membrane regions were annotated manually based on membrane predictions using the TMHMM Server v 2.0 available at http://www.cbs.dtu.dk/services/TMHMM/.

### Antibody generation

Drosophila Tmep cDNA was used as a template to generate PCR fragments corresponding to two selected peptide fragments of Tmep 1-67 (N-terminal) and Tmep 393-486 (C-terminal) using oligonucleotides (For a.a. 1-67: CF044-1F GGGGCCCCTGGGATCCATGAGCACAACCGCCGCCAG, CF044-1R GGGAATTCGGGGATCTTACTGGGCGGTCTTTGTCTGCAG for a.a 393-486: CF044-4F GGGGCCCCTGGGATCCGGTGGCAAGAATTCCCGCGGCATTC CF044-4R GGGAATTCGGGGATCTTACTGGTACTCATCGTCGCTGCTC). After amplification with KOD Hot-Start DNA polymerase (Novagen), the DNA was incubated in an InFusion (Takara Bio) reaction with *Bam*HI linearised pGEX-6p-3. The product was transformed into *E. coli* XL1-blue (Stratagene) and the sequence confirmed. The resulting constructs yielded an N-terminal fusion of GST to the CG12004 peptide sequences. The expression vector was transformed into *E. coli* Codon plus (DE3) (Stratagene) for expression studies. Overnight cultures containing LB, ampicillin (100μg/ml) were inoculated from glycerol stocks and incubated at 37°C and 200rpm. The cultures were then used to inoculate 500 ml of LB containing ampicillin (100μg/ml) before further incubation at 37°C and 200rpm. At a culture OD_600nm_ of 0.6Au, IPTG was added to a final concentration of 1 mM and incubation was continued at 18°C and 200rpm for 20 hours. The induced culture was centrifuged at 7,000*g* for 10 min at 4°C before the supernatant was discarded and the pellet was stored at −80°C until required. The purification of GST-tagged protein was performed by affinity chromatography using the following procedure. Frozen cell pellets were thawed and resuspended in lysis buffer - PBS, pH 7.2 containing protease inhibitor cocktail (Roche), prior to disruption by sonication. The supernatant was then recovered by centrifugation at 35,000*g* for 10 min at 4°C and then filtered (0.2μm). The crude extract was applied to 1ml Glutathione Sepharose beads (GE Healthcare) previously equilibrated in Buffer A (PBS) for 90min at 4°C. On completion of binding, the resin was washed three times with 50 bed-volumes (CV) of Buffer A before isocratic elution of specific protein with Buffer B (50mM Tris-HCl, pH 8.0, 10mM reduced glutathione). Fractions (500 μl) containing the GST-tagged CG12004 peptide sequences were confirmed by SDS-PAGE. Purified GST tagged TMEM184B peptides TMEM184B 1-67 and TMEM184B 393-486 in equal proportions (1:1) were injected into rabbits using the Eurogentec Polyclonal Antibody Production service (Eurogentec, Belgium) using an 87 day protocol. To affinity purify anti-Tmep antibodies, 40μg of each purified peptide were mixed, run on an SDS-PAGE gel and blotted to PVDF. The membrane was blocked with 5% milk and incubated with 1:100 anti-Tmep serum in 1% milk overnight at 4°C. The antibody was eluted using 0.2M glycine solution, pH 2.3 and neutralized with 1M Tris base.

### Viability Measurement

*Tmep^Cri^/TM3*,*Sb* females were crossed to *Df6087/TM6B*,*TbSb* (non-rescue) or *Df6087*, *UAS-mKate-Tmep/TM6B,TbSb* (rescue) males. In these crosses, because *TM3/TM6* is viable, we expected 25% of each cross to be non-Sb if the Tmep genotype was fully viable. We compared the resulting percent of non-Sb progeny to the expected for each cross. Vials in which at least 100 progeny emerged over 14 days were included in analysis.

### Locomotor Analysis

For crawling assays, wandering third instar larvae were evaluated. Larvae were rinsed quickly in water, dabbed briefly on a kimwipe to remove excess water, and placed on a clear agar dish. The number of full body pericyclic contractions were visually counted over one minute, either in real time or following video-recording and analysis. Sample sizes of 20 larvae per sex were used for every condition.

### Bang Sensitivity Analysis

Three-day-old adult flies were minimally anaesthetised with CO_2_, placed into 2.0 ml microcentrifuge tubes, and left for 10 min to recover. Flies were then vortexed for 1 min at max speed (Whirlimixer, Fisons Scientific Apparatus, U.K.). After 1 min recovery, the flies were knocked to bottom of the tube and time taken to reach the top of the tube were measured. A light was placed above the tube to promote phototaxis.

### Immunohistochemistry

Larvae were dissected in cold phosphate-buffered saline (PBS), fixed in 4% formaldehyde in PBS, washed with PBST (0.1% Triton X-100), and incubated with primary antibodies overnight at 4°C. Antibodies used include anti-Bruchpilot (Developmental Studies Hybridoma Bank (DSHB), antibody nc82, used at 1:50), anti-HRP (Jackson Immunoresearch, 123-605-021, used at 1:1000), anti-RFP (ThermoFisher # R10367, used at 1:2000), anti-Repo (DSHB, 1:20), anti-Synaptotagmin [42], anti-discs-large (DLG) (DSHB, used at 1:2000) and anti-Futsch (DSHB antibody 22C10, used at 1:20). Affinity purified rabbit anti-Tmep was used at 1:20.

### Fluorescent Microscopy

For all microscopy experiments actively behaving third instar larvae were used. Image stacks were taken on a Zeiss LSM880 inverted confocal microscope (most figures) or an Elyra structured illumination microscope (Fig 4c). Maximum intensity projections or single slices were created as noted in figure legends, exported as TIFs, and analyzed in Fiji (National Institutes of Health).

### Synapse morphology analysis

Synapse morphology analysis was carried out at muscles 6.7 hemisegment A3. Boutons were defined as distinct, spherical, anti-synaptotagmin–positive varicosity contacting the muscle. Measurements were made from images using Fiji. For branching analysis, Fiji plugin NeuronJ was used (https://imagej.net/plugins/neuronj). The extent of branching was estimated by dividing synapse length by the number of branches, where a branch was defined as any section from the synapse core which was longer than 10 μm. Satellite boutons were defined according to previous work [43–45] as small laterally branched boutons emerging from a primary axis with three or less smaller bouton sprouts. Quantification was performed and represented by dividing the number of satellite boutons by synapse branch length.

### Electrophysiology

Larvae were fillet dissected in ice-cold HL 3.1 with 0.1 mM calcium, and then washed into 0.4 mM calcium HL 3.1 for recordings of muscle 6, abdominal segment 3. A baseline recording of mEJPs was recorded for 15-30 seconds. Axons were then stimulated (3.3 mV, 0.1 ms, 0.2 Hz) at least 10 times. To be included in analysis, baseline muscle potential needed to be between - 55 and −72 mV, and baseline could not deviate more than 10 mV within this range during the entire recording. Once chosen, baselines were manually adjusted to fit a straight line using Clampex (versions 10.7 and 11.2). Using the threshold search function, we set a low threshold for detection of mEJP (0.2 mV) and EJP (5 mV) and manually excluded those signals that were above threshold but were not appropriately called. This enabled the inclusion and quantification of small multi-EJP events in mutant larvae.

### Transmission Electron Microscopy

Third instar wandering larvae were dissected and fixed in 0.1M NaPO_4_ (pH 7.4), 1% gluteraldehyde and 4 % formaldehyde (pH 7.3) overnight. Fixed larval preparations were washed 3x in 0.1 M NaPO_4_ prior to incubation in OsO_4_ (1% in 0.1 M NaPO_4_, 2 hr). Preparations were washed 3x in dH_2_O before incubation in 1% uranyl acetate. Preparations were washed (3x dH_2_O) and dehydrated through a graded ethanol series; 20 % increments starting at 30% followed by two 100 % changes, then 2x 100% propylene oxide. Preps were incubated in a graded series of Epon araldite resin (in propylene oxide); 25% increments culminating in 3x 100 % changes. Individual muscles and the ventral nerve cord were then dissected out. These were then transferred into embedding molds and the resin polymerized at 60°C for 48 hours. Resin mounted preps were sectioned (60 – 70 nm) using glass knives upon a Leica Ultracut UCT microtome and placed onto grids. Preps were subsequently incubated in uranyl acetate (50% in ethanol), washed in dH_2_O and incubated in lead citrate. Sections were imaged using a TECNAI 12 G^2^ transmission electron microscope with an SIS Megaview camera and Tecnai user interface v2.1.8 and analySIS v3.2.

